# Blunting of insulin-stimulated glucose uptake in brown adipose tissue induces systemic metabolic dysregulation in female mice

**DOI:** 10.1101/2020.12.09.416685

**Authors:** Belén Picatoste, Lucie Yammine, Rosemary Leahey, David Soares, Paul Cohen, Timothy E. McGraw

## Abstract

The role of brown adipose tissue (BAT) in thermogenesis is widely appreciated, whereas its more recently described role in whole-body metabolism is not as well understood. Here we demonstrate that deletion of Rab10 from brown adipocytes reduces insulin-stimulated glucose transport by inhibiting translocation of the GLUT4 glucose transporter to the plasma membrane. This blunting of glucose uptake into brown adipocytes induces glucose intolerance and insulin-resistance in female but not male mice. The defect in glucose uptake does not affect the thermogenic function of BAT, and the dysregulation of whole-body metabolism is independent of the thermogenic function of BAT, thereby revealing a metabolism-specific role for BAT in female mice. The reduced glucose uptake induced by RAB10 deletion disrupts ChREBP regulation of the expression of de novo lipogenesis-related (DNL) genes, providing a link between DNL in BAT and whole-body metabolic regulation that is independent of thermogenesis.

## Introduction

Insulin-stimulated glucose uptake into fat and muscle is mediated by redistribution of the glucose transporter protein 4 (**GLUT4**) from intracellular storage sites to the plasma membrane (Klip et al., 2019). Deletion of the small GTPase, Rab10, from white adipocytes blunts GLUT4 translocation and insulin-stimulated glucose uptake by about 50% (Sano et al., 2007; Vazirani et al., 2016). This partial blunting of insulin-stimulated glucose uptake by white adipocytes is sufficient to induce a systemic insulin resistance, primarily due to hepatic insulin resistance (Vazirani et al., 2016). The systemic insulin resistance of adipose Rab10 knockout mice is similar to that of adipose GLUT4 knockout, even though in the latter model there is a complete loss of insulin-stimulated glucose uptake into adipocytes (Abel et al., 2001).

The partial reduction of insulin-stimulated glucose uptake in adipose Rab10 knockout linked to a severe disruption in liver insulin sensitivity supports a metabolic sensing role for adipose-glucose metabolism. Although the exact mechanism(s) underlying adipose glucose sensing and its impact on whole body metabolic regulation have not been defined, expression of the β isoform of carbohydrate responsive-element binding protein (**ChREBPβ**) is affected in both adipose-specific GLUT4 and Rab10 KO knockout mouse models (Herman et al., 2012; Vazirani et al., 2016). ChREBP is a carbohydrate-activated transcription factor that controls key metabolic activities (Ortega-Prieto & Postic, 2019).

Previous studies documenting the role of adipose glucose metabolism in the control of whole-body metabolism relied on adiponectin-Cre to generate conditional knockout mice (Abel et al., 2001; Vazirani et al., 2016). Because adiponectin is expressed by both white and brown fat, the insulin-resistant phenotypes could be due to changes in white, brown or both types of adipocytes. Brown adipose tissue (**BAT**) has mainly been described as a thermoregulatory organ, but recent studies document a role for BAT in expending energy and in the regulation of metabolism (Blondin & Carpentier, 2016; Chondronikola et al., 2014; Cypess et al., 2015; Yoneshiro et al., 2013). Notably, insulin stimulates glucose uptake into BAT (Orava et al., 2011) and glucose accumulation in BAT is reduced in insulin-resistant individuals supporting a direct role for BAT in postprandial glucose homeostasis (Blondin et al., 2015).

We here used immortalized murine brown adipocytes to show that Rab10 deletion blunted insulin stimulated GLUT4 translocation and glucose uptake. To further unravel the role of BAT in insulin regulation of metabolism, we generated a BAT-specific RAB10 knockout mouse (**bR10KO**). Reduced glucose uptake solely into BAT induces glucose intolerance and insulin resistance in female mice. The metabolic change is independent of the role of BAT in thermogenesis. ChREBPβ expression is reduced in BAT of bR10KO mice, linking the metabolic phenotype to ChREBP-dependent glucose sensing. Despite the prominent role of BAT in metabolic regulation of female mice, there is no metabolic phenotype in male bR10KO mice.

## Results

### Rab10 silencing blunts insulin-stimulated GLUT4 translocation to the plasma membrane and glucose uptake in brown adipocytes

To study the behavior of GLUT4 in brown adipocytes, we used a dual-tagged GLUT4 reporter, HA-GLUT4-GFP, transiently expressed in brown adipocytes differentiated from immortalized murine pre-brown adipocytes (Klein et al., 2002). The ratio of the anti-HA fluorescence (surface GLUT4) to GFP fluorescence (total GLUT4) reflects the proportion of GLUT4 in the plasma membrane per cell (Lampson, Racz, Cushman, & McGraw, 2000). Insulin-stimulation induced an approximate 4-fold increase of GLUT4 in the plasma membrane of WT brown adipocytes (Figure 1A & B). Stable expression of shRNA targeting Rab10 reduced Rab10 mRNA expression by 80% (Figure 1C). This Rab10 knockdown blunted the redistribution of GLUT4 to the plasma membrane by about 50% (Figure 1A & B), with a corresponding reduction in insulin-stimulated glucose uptake, demonstrating a role for Rab10 in insulin control of GLUT4 in culture brown adipocytes (Figure D).

**Figure 1.**
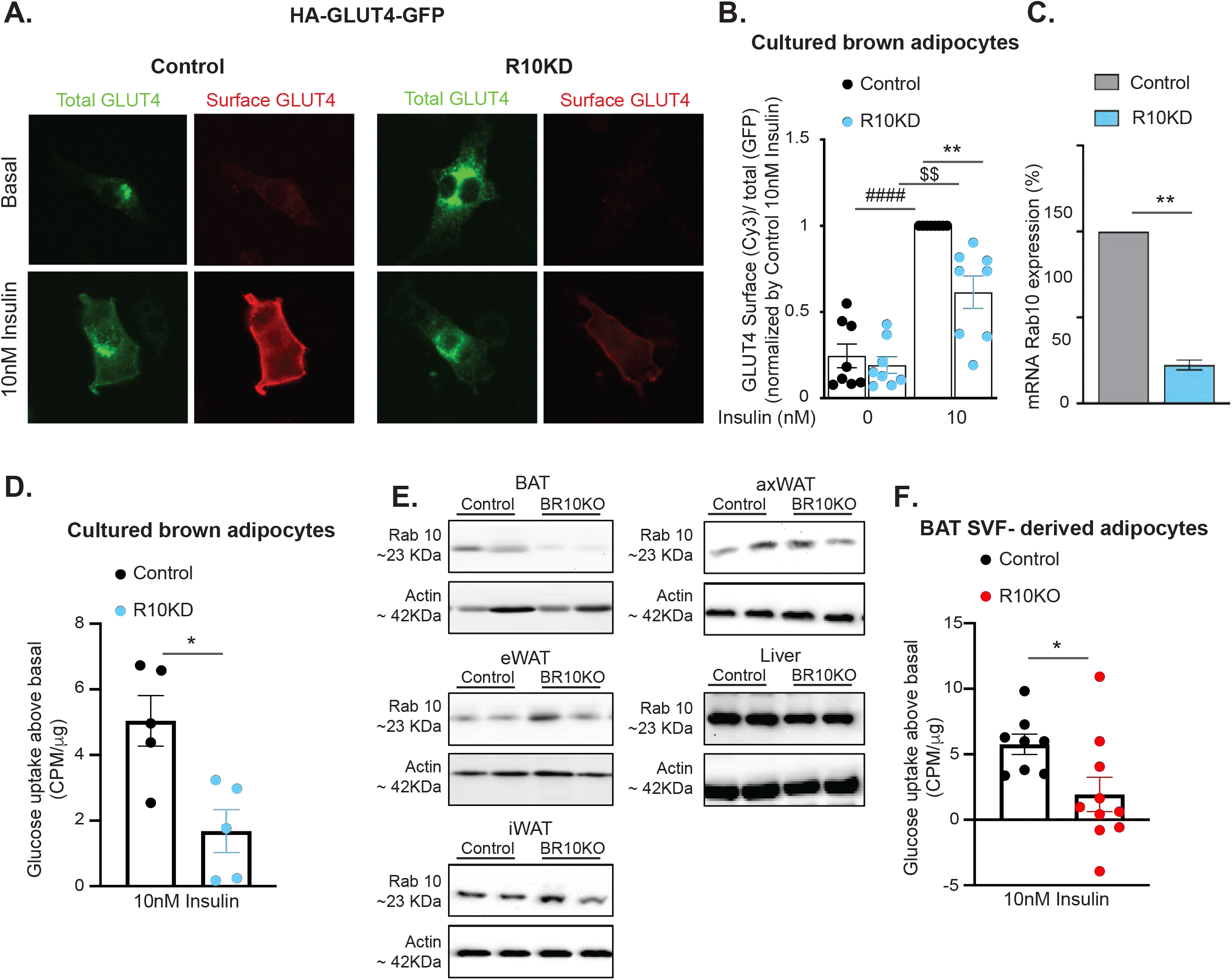
Rab10 deletion impairs insulin-stimulated GLUT4 translocatio and glucose uptake in brown adipocytes. A. Immunofluorescence staining of basal and 10nm/L insulin-treated cultured murine brown adipocytes expressing HA-GLUT4-GFP. B. Quantification of the surface-to-total ratio of HA-GLUT4-GFP in cultured murine brown adipocytes (n=8 experiments). C. Rab10 mRNA levels in control and Rab10KD cultured murine brown adipocytes (n=3 experiments). D. Insulin-stimulated glucose uptake above basal in cultured murine brown adipocytes (n=5 mice). E. Rab10 representative immunoblots of brown adipose tissue (BAT), epididymal white adipose tissue (eWAT), inguinal white adipose tissue (iWAT), axillar white adipose tissue (axWAT) and liver extracts from control and bR10KO mice. F. Insulin-stimulated glucose uptake above basal in BAT SVF-derived adipocytes (n=8-10 mice/group). *p<0.05, **p<0.001 vs control. #### p<0.0001 vs control basal. $$ p<0.001 vs R10KD basal.

To generate a model to study the role of Rab10 in primary brown adipocytes, we crossed UCP1-Cre transgenic mice to mice homozygous for loxP sites flanking exon 1 of Rab10. Rab10 protein levels were reduced in BAT-specific Rab10 knockout mouse (**bR10KO**) but unaffected in WAT depots and liver, confirming the specificity of the deletion (Figure 1E). To study the effect of Rab10 deletion in insulin-stimulated glucose uptake in brown adipocytes from our mice, we isolated BAT-stromal vascular fraction (SVF) cells that were differentiated to adipocytes (Supplementary Figure 1). Rab10 deletion blunted insulin-stimulated glucose uptake into primary BAT-SVF differentiated adipocytes (Figure 1F). These data establish, in two different models of brown adipocytes, a role for Rab10 in GLUT4 translocation and therefore in glucose uptake. These findings are in agreement with the previously documented role for Rab10 in insulin-stimulated GLUT4 translocation to the plasma membrane of white adipocytes, identifying Rab10 as essential for regulation of GLUT4 in adipocytes (Sano et al., 2007; Vazirani et al., 2016).

### Reduced glucose tolerance and systemic insulin resistance in female but not male bR10KO mice

We used bR10KO mice to explore the role of insulin-stimulated glucose uptake by brown adipocytes on whole body metabolism. Deletion of Rab10 in brown adipocytes did not significantly change the body weight of male mice fed a normal chow diet (**NCD**) (Figure 2A). Intraperitoneal glucose tolerance test (**IP-GTT**), circulating insulin levels during an IP-GTT, and insulin tolerance (**ITT**) were all unchanged in bR10KO male mice as compared to their littermate control mice (Figures 2B-D). Interestingly, adipose-specific R10KO (**aR10KO**) male mice (Rab10 deletion in WAT and BAT) were insulin resistant as compared to control and bR10KO mice (Figure 2E).

**Figure 2.**
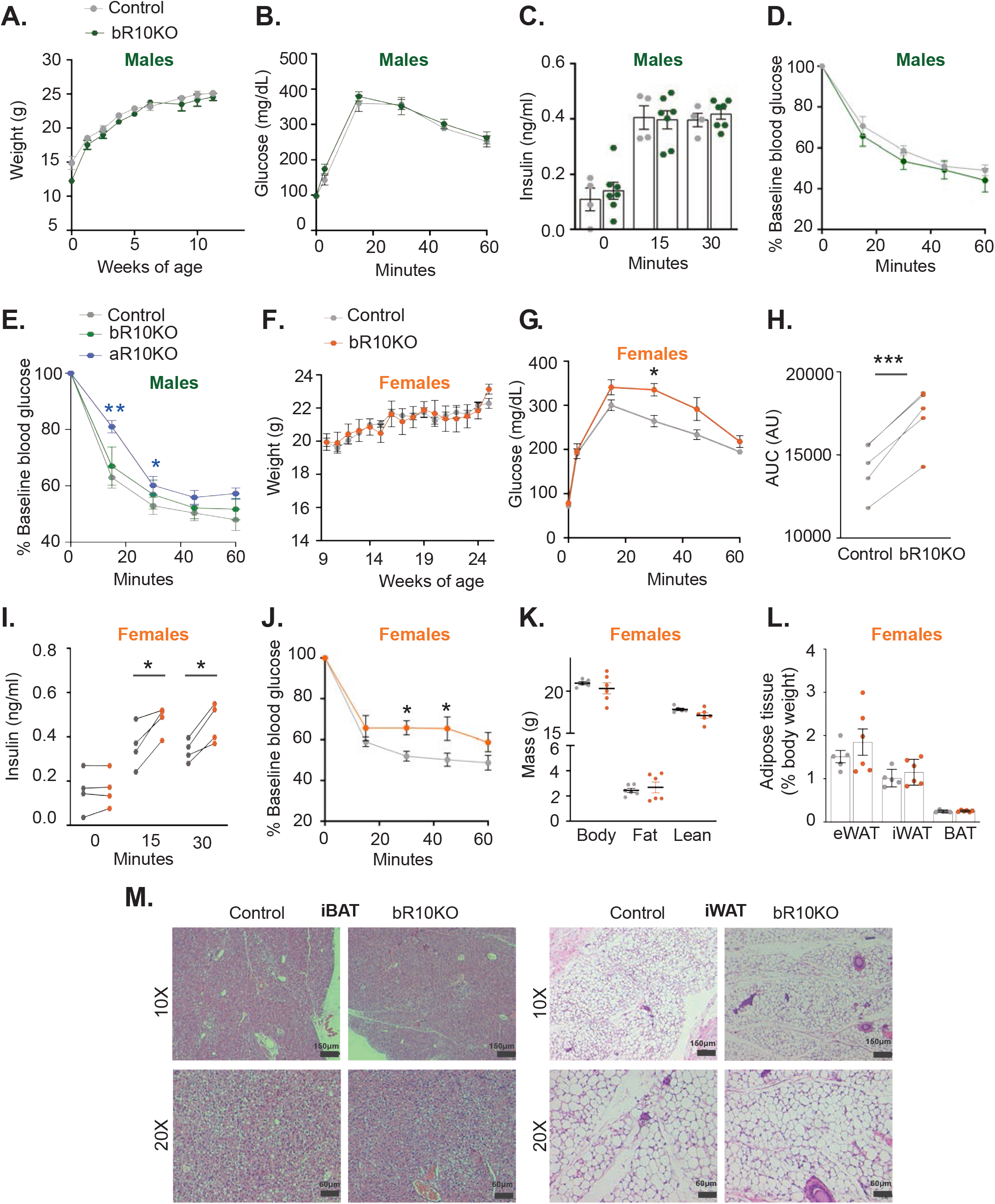
Loss of BAT-specific Rab10 is sufficient to induce systemic insulin resistance in chow diet-fed females but not males. A. Body weight overtime in male mice (*ad libitum*) (n=4-5 mice/group). B. Representative glucose tolerance test (GTT) (2g/Kg glucose intraperitoneal injection (i.p.)) in 17 weeks of age male mice (n=4-5 mice/group). C. Plasma insulin levels in male mice after overnight fasting (0 minutes) and 15 and 30 minutes after i.p. glucose injection (2g/Kg) (n=4-7 mice/group). D. Representative insulin tolerance test (ITT) results for 17 weeks of age (0.75 unit/Kg insulin i.p.) (n=4-6 mice/group). E. Representative insulin tolerance test (ITT) results for 16 weeks of age (0.75 unit/Kg insulin i.p.) (n=4-6 mice/group). F. Body weight overtime in female mice (*ad libitum*) (n=4-5 mice/group). G. Representative glucose tolerance test (GTT) (2g/Kg glucose intraperitoneal injection (i.p.)) in 17 weeks of age female mice (n=4-5 mice/group). H. Quantification of GTT area under the curve (averages of n=5 independent cohorts of mice of 17-20 weeks of age. I. Plasma insulin levels in female mice after overnight fasting (0 minutes) and 15 and 30 minutes after i.p. glucose injection (2g/Kg) (averages for n=4 independent cohorts of mice of 17-20 weeks of age). J. Representative ITT results for 17 weeks of age (0.75 unit/Kg insulin i.p). K. Body composition by magnetic resonance imaging (MRI) in 12 weeks of age mice (n=6 mice/group). L.Adipose tissue depots weights: epididymal adipose tissue (eWAT), inguinal adipose tissue (iWAT) and brown adipose tissue (BAT) in 25 weeks of age mice (n=5-6 mice/group). M. Representative H&E images of BAT and iWAT from control and bR10KO female mice. Images are shown at 10X and 20X magnification with 150μm and 60μm scale bars respectively. *p< 0.05, **p< 0.01, ***p<0.0005.

Similarly, deletion of Rab10 in brown adipocytes did not affect body weight of female mice on NCD (Figure 2F). However, bR10KO female mice were glucose intolerant with elevated plasma insulin during the IP-GTT (Figures 2G-I). In agreement with the elevated blood insulin, bR10KO female mice were insulin resistant in an ITT (Figure 2J). Despite this impact of Rab10 deletion in brown adipocytes on metabolism of female mice, there were no differences in whole-body fat mass, the weights of individual fat depots and overall fat tissue anatomy between genotypes of female mice (Figure 2K-M).

### Brown fat Rab10 knockout does not augment the metabolic impacts of a high fat diet

To explore the relationship between the altered BAT glucose metabolism caused by Rab10 deletion and the metabolic insult of over-nutrition, we challenged bR10KO mice with high fat diet (HFD: 20% protein, 60% fat, 20% carbohydrate). Deletion of Rab10 from brown adipocytes did not affect weight gain or WAT and BAT fat mass, in female or male mice on a HFD (Figures 3A-D). In addition, there were no differences within the sexes between HFD fed bR10KO and control mice in glucose tolerance, plasma insulin during a GTT and insulin sensitivity (Figures 3E-L). Thus, disruption of brown fat metabolism induced by Rab10 KO does not have additive effects on the impacts of HFD on whole-body metabolism.

**Figure 3.**
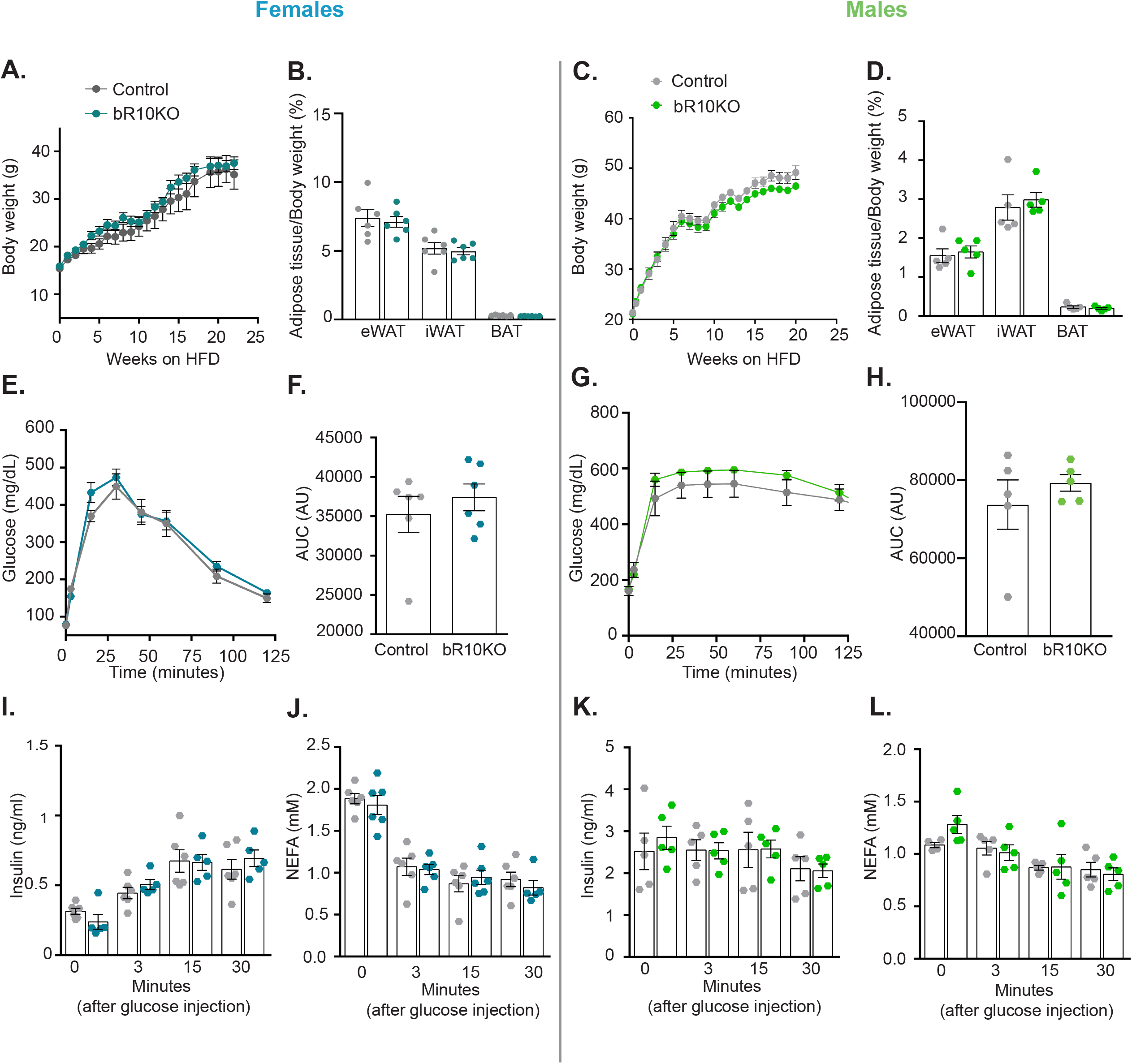
BAT Rab10 knockout does not augment the metabolic impacts of a high fat diet. Female and male mice were ad libitum fed HFD starting at 6 weeks of age. A. Female mice body weight (ad libitum) through 17 weeks of a HFD. B. Adipose tissue depots weights: epididymal adipose tissue (eWAT), inguinal adipose tissue (iWAT) and brown adipose tissue (BAT) of female mice after 20 weeks of a HFD. C. Male mice body weight (ad libitum) through 25 weeks of a HFD. D. Adipose tissue depots weights: eWAT, iWAT and BAT of male mice after 25 weeks of a HFD. E. Glucose tolerance test (GTT) (2g/Kg glucose intraperitoneal injection (i.p.)) in 13 weeks of age female mice. F. Quantification of GTT area under the curve (AUC). G.GTT (2g/Kg glucose i.p.) in 17 weeks of age male mice. H. Quantification of GTT area under the curve (AUC). I. Plasma insulin and J. free fatty acid levels in female mice after overnight fasting (time point 0) and 3, 15 and 30 minutes after i.p. glucose injection (2g/Kg). K. Plasma insulin and L. free fatty acid levels in male mice after overnight fasting (0 minutes) and 3, 15 and 30 minutes after i.p. glucose injection (2g/Kg). Female mice data n=6 mice/group and male mice data n=5 mice/group.

### BAT-specific Rab10 deletion is not required to maintain body temperature with cold exposure

To test whether Rab10 deletion interferes with BAT’s function in thermogenesis, bR10KO and control male and female mice were adapted to severe cold (4°C) for 2 weeks. Body temperature was measured every 30 minutes for the first 3 hours and thereafter once a day for 14 days (Figures 4A-D). Body temperature did not differ between control and bR10KO mice in both sexes, demonstrating that BAT thermoregulatory capacity is maintained in bR10KO mice. Body weight and adiposity did not differ between control and bR10KO animals in both sexes adapted to severe cold (Figures 4E-H), although there was a minor, albeit statistically significant, decrease in BAT weight in bR10KO female mice as compared their littermate controls (Figure 4F). Consistent with the deletion of Rab10 not affecting BAT’s role in thermogenesis, there were no major changes in the expressions of thermoregulatory genes in the bR10KO BAT (Figure 4I). In addition, there were no differences in the morphology of BAT between genotypes at either room temperature or with severe cold challenge (Figure 4J and Supplementary Figure 2).

**Figure 4.**
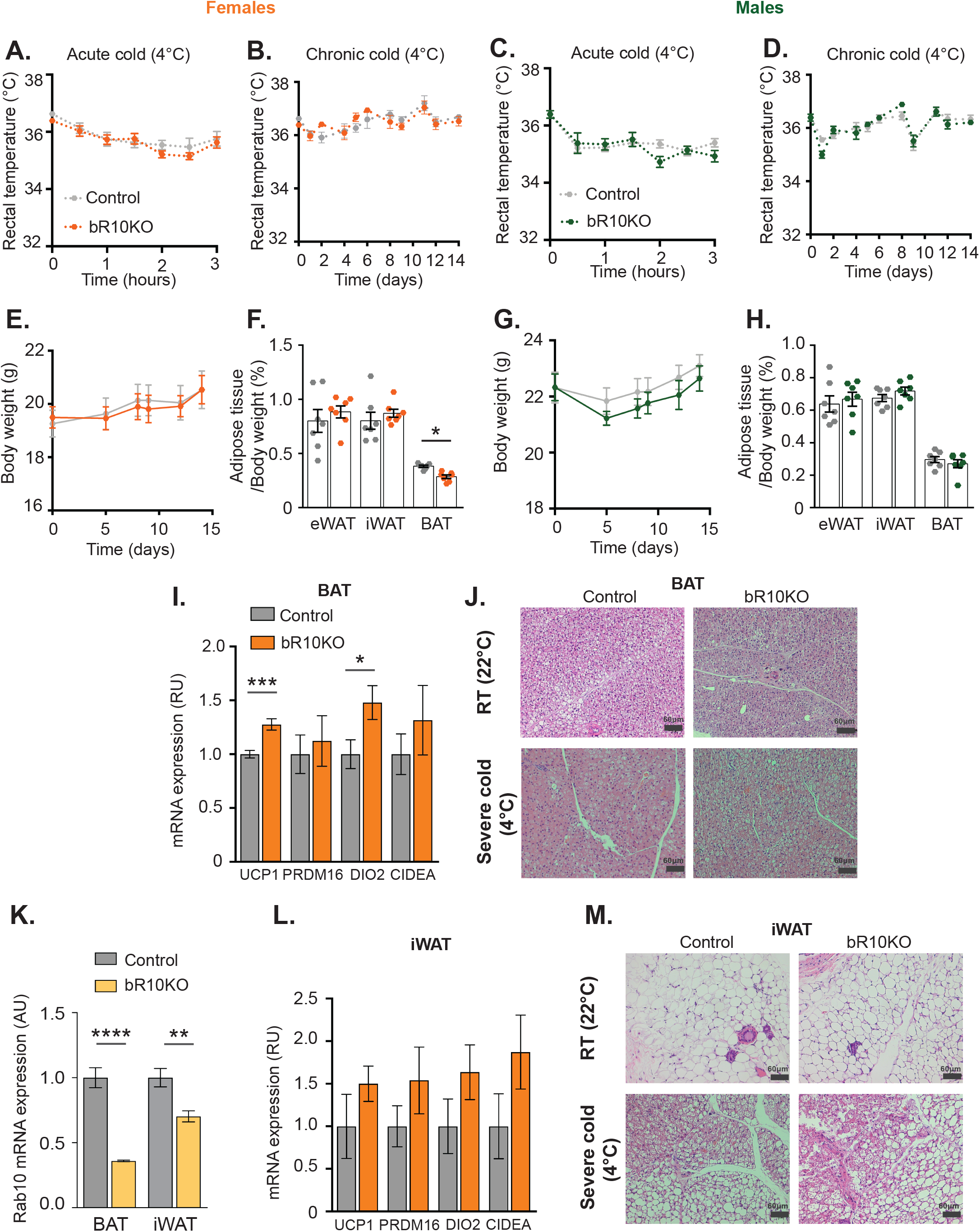
BAT Rab10 is not required to maintain body weight temperature when exposed to severe cold. A. Rectal temperature in female mice under acute cold (4°C) or B. chronic cold (4°C). C. Rectal temperature in male mice under acute cold (4°C) or B. chronic cold (4°C). E. Body weight overtime (*ad libitum*) and F. Adipose tissue depots weights: epididymal adipose tissue (eWAT), inguinal adipose tissue (iWAT) and brown adipose tissue (BAT) of female mice after 14 days at 4°C. G. Body weight overtime (*ad libitum*) and H. eWAT, iWAT and BAT weight of male mice after 14 days at 4°C. I. mRNA expression for UCP1, PRDM16, DIO and CIDEA in BAT samples of female mice after 14 days at 4°C. J. Representative H&E staining images of BAT of mice housed at room temperature (22°C) and chronic cold (4°C) conditions. Images are shown at 20X magnification with 60μm scale bars. K. mRNA expression of Rab10 in BAT samples of female mice after 14 days at 4°C. L. mRNA expression of UCP1, PRDM16, CIDEA and DIO2 in iWAT samples of female mice after 14 days at 4°C. M. Representative H&E staining images of iWAT of mice housed at room temperature (22°C) and chronic cold (4°C) conditions. Images are shown at 20X magnification with 60μm scale bars. N=7 mice/group for all the conditions and experiments. *p <0.05, **p<0.01, ***p<0.001 and ****p<0.0001.

Cold challenge induces a thermogenic gene program in subcutaneous WAT, a process referred to as “browning”, that includes the increased expression of UCP1 (Herz & Kiefer, 2019). There was a significant decrease of Rab10 mRNA levels in the iWAT from severe cold challenged bR10KO female mice, demonstrating the induction of UCP-1 driven Cre and silencing of Rab10 in the iWAT of these mice (Figure 4K). However, “browning” genes in iWAT of severe cold challenged bR10KO females were not reduced as compared to iWAT from severe cold challenged control female mice (Figure 4L). The morphologies of iWAT in both genotypes of mice were similarly affected by severe cold, characterized by a decrease in size of white adipocytes and an increase in multilocular brown-like adipocytes (Figure 4M and Supplementary Figure 2). Thus, iWAT from bR10KO female mice shows a normal response to cold exposure demonstrating that Rab10 does not have a role in WAT browning.

### Thermoneutrality-adapted bR10KO mice are insulin resistant and have increased adiposity

At thermoneutrality (30°C for mice) loss of body heat is minimal and energy consumption for regulatory thermogenesis against cold is suppressed (Lichtenbelt, Kingma, van der Lans, & Schellen, 2014). To investigate the effect of Rab10 silencing on non-thermogenic functions of BAT, male and female mice were studied after adaptation to thermoneutrality for 21 days. UCP1 expression levels in BAT decrease when mice are acclimated to 30°C. bR10KO male and female mice show a 50% reduction of Rab10 expression in BAT as compared to their wild type littermate controls (Rab10^fl/fl^) (Figure 5A).

**Figure 5.**
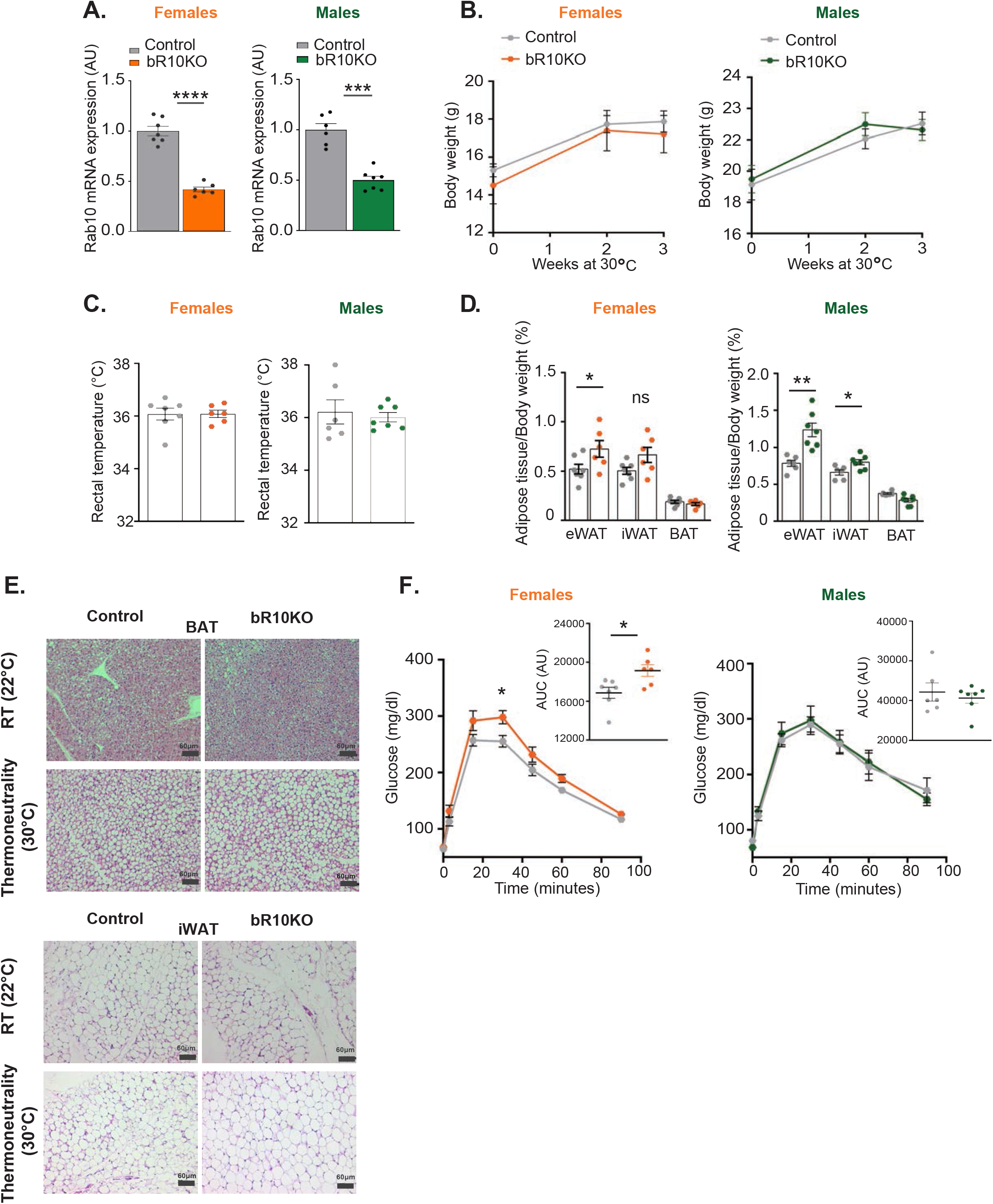
Thermoneutrality-adapted BRab10KO female mice are insulin resistant and both females and males have increased adiposity. A. Rab10 mRNA levels in BAT of female and male mice after 21 days at 30°C. B. Body weight overtime (ad libitum) of female and male mice at 30°C. C. Rectal temperature of females and males acclimated for 3 weeks at 30°C. D. Adipose tissue depots weights: epididymal adipose tissue (eWAT), inguinal adipose tissue (iWAT) and brown adipose tissue (BAT) of female and male mice after 3 weeks at 30°C. E. Representative H&E staining images of BAT and iWAT from females acclimated to 22°C or 30°C for 3 weeks. Images are shown at 20X magnification with 60μm scale bars. F. Glucose tolerance test (GTT) (2g/kg glucose intraperitoneal injection (i.p.)) at day 19 of thermoneutrality acclimation in female and male mice. Quantification of GTT area under the curve (AUC). N=6-7 mice/group for all the conditions and experiments. *p <0.05, **p<0.01, ***p<0.001 and ****p<0.0001.

Body weight and temperature did not differ between genotypes in both sexes at thermoneutrality (Figure 5B and C) although eWAT and iWAT, as a percentage of body weight, were increased in bR10KO male and female mice (Figure 5D). Consistent with previous reports (Cui et al., 2016), there was an increase in BAT adipocyte size in control mice housed at 30°C, and no differences between genotypes were found (Figure E and Supplementary Figure 2).

Female bR10KO mice were glucose intolerant at 30°C (~15% difference in AUC), demonstrating a thermogenic-independent function of BAT in regulation of systemic metabolism that is disrupted by Rab10 deletion in female mice (Figure 5F). There were no differences in glucose tolerance between the genotypes of the male mice, re-enforcing the sex-dependence of the impact of Rab10 KO in BAT on metabolic phenotype (Figure 5F).

### ChREBPβ expression is reduced in Rab10KO BAT

ChREBPβ and *de novo* lipogenesis-related genes are induced in BAT by cold (Joan Sanchez-Gurmaches et al., 2018). ChREBPβ expression was reduced in female bR10KO BAT at thermoneutrality, mild cold stress (room temperature) and severe cold stress (Fig 6A, C and E). The mRNA and protein levels of *Acc1* and *FASN*, two proteins key for *de novo* lipogenesis (DNL), were also reduced in bR10KO BAT at all temperatures (Fig 6A, C, E and G). However, the expression of neither ChREBPα, which regulates ChREBPβ expression (Herman, 2012), nor *Srebp1c*, a transcription factor involved in lipogenesis (Crewe et al, 2019), were affected by Rab10 KO (Fig 6B, D and F). Thus, the blunting of insulin-regulated glucose uptake induced by Rab10 deletion disrupts the expression in BAT of genes that control lipid metabolism independent of thermoregulation of these genes.

**Figure 6.**
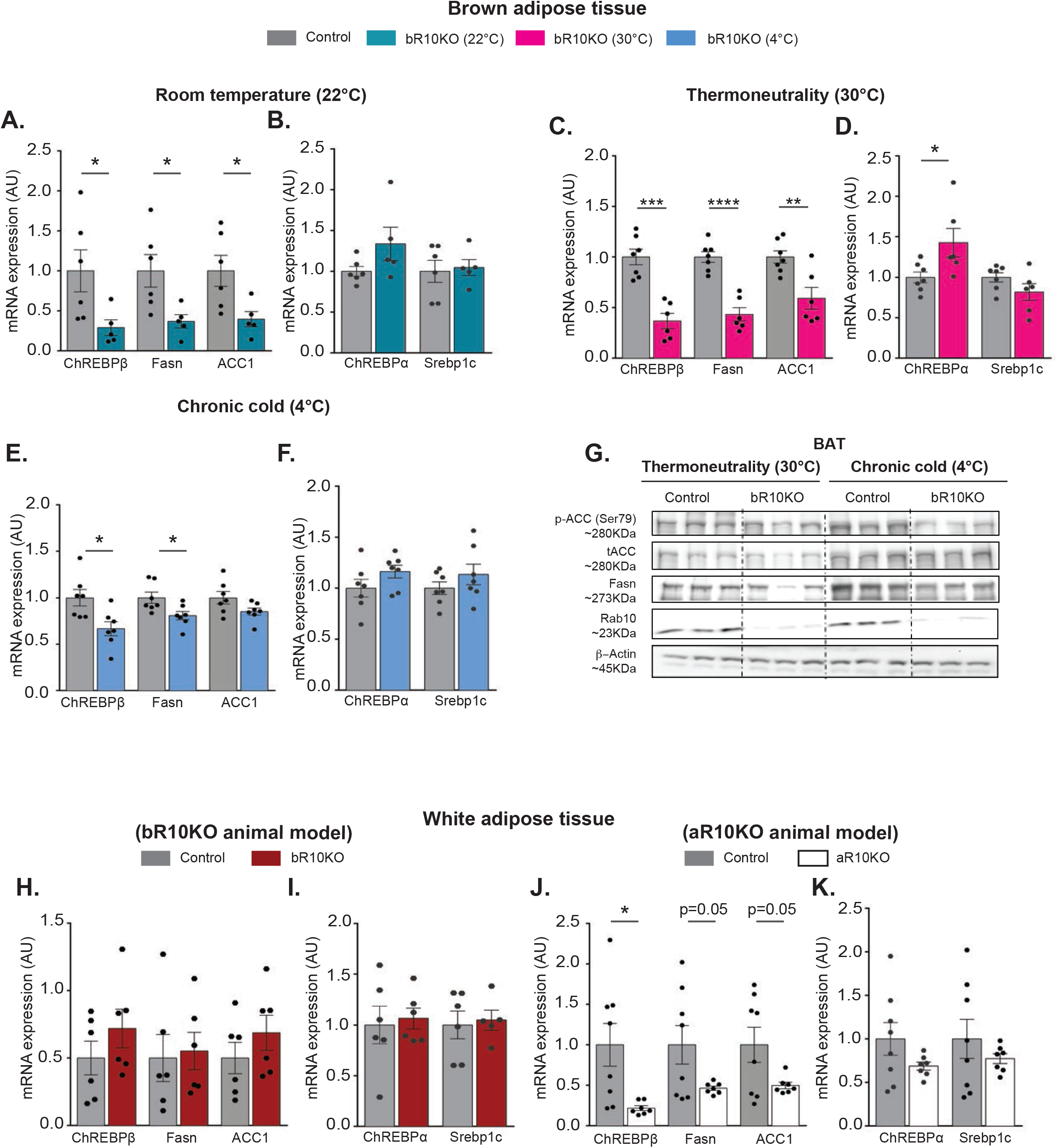
ChREBPβ and DNL-related gene expression are blunted in Rab10KO BAT and eWAT in a cell-intrinsic manner. mRNA expression for ChREBPβ, FASN, ACC, ChREBPα and Srebp1c in BAT samples from 9-12 weeks of age female mice: A-B. At room temperature or mild cold (22°C) (n=5-6 mice/group), C-D. At thermoneutrality (30°C) for 3 weeks (n=6-7 mice/group) and E-F at severe cold (4°C) for 14 days (n=7 mice/group). G. A. Representative immunoblots of brown adipose tissue (BAT) extracts from control and bR10KO mice at 30°C and 4°C. H-K. mRNA expression for ChREBPβ, FASN, ACC1, ChREBPα and Srebp1c in eWAT samples from G-H. 12 weeks of age control and bR10KO females. *p<0.05, **p<0.01***p<0.001 and ****p<0.0001.

ChREBPβ, *Acc1* and *FASN* mRNA levels were not reduced in eWAT from bR10KO mice demonstrating that the reduction is a cell-intrinsic effect of RAB10 deletion (Fig 6H-I). Consistent with a cell-intrinsic effect of Rab10 deletion on the expression of DNL genes, Rab10 silencing in eWAT induces reduced expression of DNL genes (Fig 6J-K). Furthermore, expression of DNL genes was reduced in BAT from male bR10KO mice, reinforcing the link between reduced insulin-stimulated glucose transport induced by Rab10 deletion with reduced expression of DNL genes (Supplementary figure 3A and B).

## Discussion

Glucose uptake by adipose tissue accounts for a minor fraction (10 to 15%) of a prandial glucose load (Kahn, 1996). Nonetheless, previous studies of mouse models with alterations in adipose insulin-regulated glucose metabolism have revealed the essential contributions of adipose glucose metabolism to systemic metabolism (Abel et al., 2001; Blüher et al., 2002; Tanaka et al., 2015; Vazirani et al., 2016; Vijayakumar et al., 2017; Skorobogatko et al., 2018). In the majority of those studies WAT was the primary focus for understanding the link between adipose glucose metabolism and whole-body metabolic alterations. However, the adiponectin promoter driven CRE used to generate the knockout models deletes genes from both white and brown adipocytes. Therefore, the whole-body metabolic dysregulation could be due to changes in both WAT and BAT. Here we undertook a study of reduced insulin-stimulated glucose uptake in brown adipocytes induced by knockout of Rab10, a component of the machinery linking insulin receptor activation to GLUT4 translocation in white adipocytes. Our data reveal a role for Rab10 in the regulation of GLUT4 in brown adipocytes and this disruption in BAT metabolism leads to whole-body insulin resistance in female mice without affecting thermoregulation. Brown fat specific knockout of Rab10 did not result in whole-body metabolic disruptions in male mice. These results contribute to an enhance understanding of the role of brown adipocytes in metabolic regulation independent of thermoregulation.

### Rab10 is required for GLUT4 translocation in brown and white adipocytes

It has previously been established that Rab10 is required for insulin-stimulated GLUT4 translocation in white adipocytes, and here we demonstrate that Rab10 is also required for GLUT4 translocation in brown adipocytes. Brown adipocytes originate from a common precursor shared with skeletal muscle (Seale et al., 2008; J. Sanchez-Gurmaches & Guertin., 2014). Studies of cultured rat muscle cells have shown that the Rab8A but not Rab10 controls muscle GLUT4 traffic in response to insulin (Sun, Bilan, Liu, & Klip., 2010). Thus, despite the common lineage between brown adipocytes and muscle cells, adipocytes share components of the machinery that regulates GLUT4 traffic and glucose uptake which are distinct from regulation in muscle.

### Regulated glucose uptake contributes to BAT sensing nutritional states in female mice

Prior studies have established that reduced BAT mass and/or activity are associated with metabolic syndrome and obesity (Bartelt et al., 2011; Stanford et al., 2013). Here, we extend the understanding of BAT as a metabolic tissue by demonstrating that a partial reduction in GLUT4 translocation in brown adipocytes is sufficient to induce insulin resistance in female mice. The reduced glucose uptake is not associated with changes in BAT mass or the size of brown adipocytes, indicating that anabolic activities in Rab10 KO brown adipocytes are intact. Our results suggest that insulin-regulated glucose uptake by brown adipocytes has a role in metabolic sensing and that the approximate 50% reduction in postprandial glucose uptake resulting from blunted GLUT4 translocation is sufficient to disrupt this sensing. Importantly, Rab10 deletion in BAT affects whole-body insulin sensitivity at thermoneutrality as well as at mild cold stress, uncoupling thermoregulation of BAT function from BAT regulation of metabolism. Furthermore, our findings support a role for BAT dysfunction contributing to the natural development of insulin-resistance since they demonstrate that even partial disruptions in BAT glucose metabolism, as might occur in the early stages of development of disease, have an outsize effect on whole body metabolism.

In agreement with a role in metabolic sensing for glucose metabolism in brown adipocytes, enhanced glucose flux into brown adipocytes, induced by knockout of RalGAPB, a negative regulator of GLUT4 translocation to the plasma membrane, is associated with enhanced glucose tolerance (Leto et al., 2012; Chen et al., 2007; Skorobogatko et al., 2018). Thus, brown adipocytes have a rheostat function in metabolic homeostasis, where decreased brown adipocyte glucose metabolism is associated with reduced whole-body insulin-sensitivity, whereas enhanced brown adipocyte glucose metabolism is associated with improved insulin-sensitivity.

Unlike female mice, bR10KO male mice fed a NCD have normal glucose tolerance and insulin sensitivity. Differences between male and female mice in cold- or diet-induced thermogenesis have been previously documented, consistent with sex differences in BAT function in thermogenesis (Quevedo, Roca, Picó, & Palou, 1998; Roca et al., 1999). Our data extend differences between sexes in BAT function to include the impact of BAT on systemic insulin sensitivity. The reasons for differences in the impact of Rab10 deletion from brown adipocytes between male and female mice will be explored in future studies.

### CHREBPβ a glucose sensor in brown adipocytes

ChREBP is a glucose-sensing unit critical for controlling insulin sensitivity in mice and in humans (Baraille, Planchais, Dentin, Guilmeau, & Postic, 2015; Herman et al., 2012). Insulin resistance is associated with lower ChREBP and DNL-related gene expression in WAT of humans and rodents (Eissing et al., 2013; Herman et al., 2012; Kursawe et al., 2013; Roberts et al., 2009). ChREBP regulation involves direct activation of ChREBPα in response to high glucose/fructose that activates expression of ChREBPβ, which in turn activates transcription of DNL-related genes (Herman et al., 2012). The reduced expression of ChREBPβ, ACC1 and FASN in Rab10KO BAT female mice support ChREBP being a mediator of the Rab10 knockout effect. Consistent with a link between Rab10 knockout-induced changes in glucose uptake and ChREBP activity, the effect of Rab10 deletion on ChREBPβ expression was cell intrinsic. Expression of ChREBPβ and DNL-related genes in both BAT and WAT of aR10KO female mice (Rab10 deletion in WAT and BAT) were reduced, whereas the expression of these genes were unaffected in WAT of bR10KO female mice (Rab10 deletion solely in BAT).

The expression of ChREBPβ and DNL genes in WAT and BAT of male mice were also reduced in a cell intrinsic manner by Rab10 knockout, re-enforcing the link between Rab10 knockout and reduced ChREBP activation. However, in male mice, only when Rab10 was silenced in WAT (adiponectin-CRE) was the loss of Rab10 linked to insulin resistance, reenforcing the dominant role of WAT in metabolic control of male mice.

Previous studies show that in *ad lib-fed* wild type mice, the rate of *de novo* lipogenesis (DNL) in BAT is ~40-fold higher than in WAT and that adipose-ChREBP is required for induction of this process in both WAT and BAT in response to sucrose (Vijayakumar et al., 2017). Our results are in line with the silencing of ChREBP in adipocytes causing systemic insulin resistance (Vijayakumar et al., 2017). However, different genetic murine models manipulating DNL pathway in adipocytes have reported inconsistent results regarding systemic effects (Vijayakumar et al., 2017; Herman et al., 2012; Lodhi et al., 2012; Carvalho et al., 2005). For example, a study of a tamoxifen-inducible adipose-specific FASN knockout, which blunts DNL in adipocytes, showed improved systemic glucose tolerance (Guilherme et al., 2017). An explanation for the differences in results is that disruption of the ChREBP pathway might impact the synthesis of specific lipids and or perturb the balance of lipid species in ways distinct from the loss of FASN. It has recently been shown that the effect of loss of adipocyte FASN is in part mediated through M2 macrophages and promotion of beiging of WAT (Henriques, 2020). In our brown adipocyte Rab10 knockout model we do not see an increase of beige adipocytes in WAT, supporting the hypothesis that disruption of ChREBP pathway is functionally distinct from the loss of FASN.

Circulating exogenous fatty acids and glucose provide fuel for catabolism by BAT during prolonged cold (Bartelt et al., 2011; Labbé et al., 2015). It has also been shown that BAT lipogenesis induced by cold challenge is required for optimal thermogenesis and fuel storage, processes linked to AKT2 activity (Sanchez-Gurmaches et al., 2018). AKT2 is also required for insulin-stimulated glucose uptake (González & McGraw, 2009); however, the effect of AKT2 on DNL in brown adipocytes is not rescued by increases in glucose uptake alone, suggesting AKT2 regulated activities, in addition to enhanced glucose uptake, are required for stimulation of DNL (Sanchez-Gurmaches et al., 2018). In our Rab10 knockout model, glucose transport is reduced due to decreased GLUT4 translocation without an effect of AKT activation (Vazirani R et al., 2016) supporting a direct role for glucose regulation.

### BAT contribution to whole-body metabolic regulation is independent of its role in thermogenesis

Glucose oxidative metabolism in BAT contributes to the fueling of non-shivering thermogenesis during cold exposure (Townsend & Tseng, 2014). bR10KO female mice thermoregulate properly, demonstrating that the deletion of Rab10, despite disrupting wholebody metabolism, does not affect BAT’s contribution to thermal regulation. Similarly, the reduced insulin-stimulated glucose uptake in BAT-specific AKT2 KO mice is not linked to cold intolerance (Joan Sanchez-Gurmaches et al., 2018). bR10KO female mice acclimated to 30°C, a temperature at which BAT thermogenic activity is minimal, remain glucose intolerant and insulin resistant, providing additional evidence that the metabolic regulatory function of BAT is independent of thermogenesis. The uncoupling of BAT role in thermogenesis from its role in the regulation of metabolism revealed by the mouse studies are in line with human studies demonstrating normal non-shivering thermogenesis in insulin-resistant individuals despite reduced insulin-stimulated glucose uptake into BAT (Blondin et al., 2015). The crosstalk between BAT and other organs influences systemic metabolic fluxes and endocrine networks and that would be a potential explanation for the bR10KO metabolic phenotype. In this regard, BAT is reported to secrete hormones (regulatory molecules) that impact systemic metabolism (Gunawardana et al., 2012; Payab et al., 2020).

We demonstrate for the first time the role of Rab10 on insulin-stimulated GLUT4 translocation to the plasma membrane in brown adipocytes and we provide a unique model to understand new mechanisms of BAT function in controlling whole body glucose metabolism. Although there is strong support for the role of BAT DNL in controlling systemic insulin sensitivity, the molecular mechanism underlying this regulation remain to be discovered.

## Acknowledgments

We thank the metabolic phenotyping center at Weill Cornell Medicine, Marissa Cortopassi, Hayley Nicholls and Dr. David Cohen for their assistance with metabolic experiments and discussion. We thank Michele Alves-Bezerra and Mariana Acuna Aravena for their assistance with metabolic and in vitro experiments. We thank the Weill Cornell Medicine Biochemistry Microscopy & Image Analysis Core and the Weill Cornell Medicine Laboratory of Comparative Pathology for their assistance with the histology work. We thank members of the McGraw lab for critical reading of the manuscript. We thank Matthew Potthoff and Johannes Klein for the pre-brown adipocyte cells. This work was supported by NIH grants DK52852, DK096925 and DK125699 (T.E.M). MBPB is supported by a Postdoctoral Fellowship from the American Diabetes Association (grant 1-19-PMF-026).

## Author Contributions

B.P. designed and conducted experiments, analyzed the data, prepared figures and wrote the manuscript. L.Y. conducted experiments, analyzed the data and edited the manuscript. R.L. conducted experiments, analyzed the data and edited the manuscript. D.S. assisted with animal studies. P.C. gave advice and designed experiments. T.E.M. conceived and supervised the project, designed experiments, analyzed the data and wrote the manuscript.

## Declaration of Interests

The authors declare that there is no conflict of interest.

---

**Supplementary Figure 1. BAT-SVF adipocytes.** A. Immunofluorescence staining for perilipin in BAT-SVF adipocytes. B. qPCR analysis of mRNA levels of brown adipocyte markers of BAT-derived SVF cells before and after differentiation into adipocytes. *p<0.05, p<0.001, p<0.0001.

**Supplementary Figure 2**. Brown (BAT) and subcutaneous inguinal white adipose tissue (iWAT) from female mice. A. BAT and iWAT from female mice acclimated to thermoneutrality for 21 days. B. BAT and iWAT from female mice housed at room temperature. C. BAT and iWAT from female mice acclimated to severe cold for 14 days. Images are shown at 10X and 20X magnification with 50μm and 20μm scale bars respectively.

**Supplementary Figure 3. ChREBPβ and DNL-related gene expression are blunted in Rab10KO BAT from male mice.** mRNA expression for A. ChREBPβ, FASN and ACC and B. ChREBPα and Srebp1c in BAT samples from 10 weeks of age thermoneutrality-acclimated male mice. *p <0.05, ***p<0.001 and ****p<0.0001.

## Research Design and Methods

### Brown adipose tissue-specific Rab10 knockout mice

Adipose-specific Rab10KO mice (aR10KO) were generated as previously published by our group (Vazirani et al., 2016). We use mice with C57BL/6J genetic background. Cre-lox recombination was used to generate the BAT-specific Rab10KO mice (bR10KO). Mice with loxP sites flanking exon 1 of Rab10 were bred with UCP-1-cre transgenic mice to specifically drive deletion of Rab10 in mature adipocytes expressing UCP1, that are mostly brown adipocytes. Heterozygotes with (Rab10^fl/wt^+ cre) and without (Rab10^fl/wt^) the UCP1-cre transgene were bred to generate mice for further breeding. Homozygote flox females (Rab10^fl/fl^ + cre) and homozygote (Rab10^fl/fl^) were bred to generate mice for the experiments. Control data were derived from Rab10^fl/fl^ littermate mice. Mice were maintained on a standard 12-h light/dark cycle and had *ad libitum* access to water and food. Mice were fed standard rodent chow diet (5053, PicoLab^®^ Rodent Diet), or 60% irradiated high fat diet (D12492, Research Diets).

Animals were handled in accordance with guidelines of the Weill Cornell Medicine Institutional Animal Care and Use Committees.

### Genotyping

Toe or tail DNA was retrieved through digestion with 50 mM NaOH & Tris and used in touchdown PCR. Cre and IRS recombination was verified by using the following primers: CRE (5’-ATGTCCAATTTACTGACC-3’ and 3’-CGCCGCATAACCAGTGAAAC-5’) and IRS (5’-GTCTTGCTCAGCCTCGCTAT-3’ and 3’-ACAGCGTGAATTTTGGAGTCAGAA-5’). Rab10 recombination was verified by using the following pair of primers, RAB10 (5’ – GGTAAAGGCAAGTAGATGTCCATG and 3’ – GAAGAGCAATTAAACACTGCATGC).

### Cell lines and culture

Immortalized murine pre-brown adipocytes (Klein et al., 2002) were cultured in Dulbecco’s Modified Eagle’s Medium - high glucose (D5796, Sigma-Aldrich) supplemented with 20% FBS (26140-095, Life Technologies), and Penicillin/Streptomycin (15070-063, Life Technologies) at 37°C/ 5% CO2. To generate the stable Rab10 KD pre-brown adipocyte cell line, we used pSIREN RetroQ system from Clontech, encoding sequences that target Rab10.

Cells were passaged at 80% confluence. For differentiation, cells were plated at approximately 60-80% confluence and differentiated with growth media containing 20nM insulin (I5500, Sigma-Aldrich) and 1nM T3 (T5516, Sigma-Aldrich) (day1). On day 3 cells were induced with growth media containing 20nM insulin, 5.1μM Dexamethasone (D4902, Sigma-Aldrich), 0.5mM IBMX (I5879, Sigma-Aldrich), and 0.125mM Indomethacin (I7378, Sigma-Aldrich). The stock solution of Indomethacin was heated for at least 4 minutes at 75°C to get into solution before adding to induction media. On day 4 cells were differentiated with the above cocktail. Experiments were performed on day 6 after differentiation.

The stromal vascular fraction (SVF) was isolated from digested iBAT (digestion buffer: 125mM NaCl, 5mM KCl, 1.3mM CaCl_2_, 5mM glucose 1% pen/strep, 100mM HEPES, 4% BSA, and 1.5mg/ml Collagenase B) and cultured in DMEM/F12 GlutaMAX (1X) media (10565018, Gibco) containing 10% FBS and 1% pen/strep. Upon confluence, cells were differentiated in DMEM/F12 media containing 10% FBS, 1 μM rosiglitazone, 1 μM Dexamethasone, 0.5mM IBMX, 17nM insulin and 1nM Thyroid hormone T3 (day 0) 1 μM rosiglitazone. After 2 days, SVF adipocytes were maintained in media containing 10% FBS, 17nM/L insulin and 1 μM rosiglitazone. Two days later, rosiglitazone was removed but insulin was continued. Experiments were performed on day 6 after differentiation.

### Glucose uptake in murine brown adipocytes and in brown adipose tissue Stromal Vascular Fraction Adipocytes

For glucose uptake experiments, cells were starved for 1 hour. Then cells were incubated in pre-warmed 0.1%BSA KRP buffer (1.2M NaCl, 48mM KCl, 12mM KH2PO4, 6mM MgSO4(H2O)7, 12mM CaCl_2_, 20mM HEPES, 100nM adenosine and 2.5% Insulin and fatty acids-free BSA) for 1 hour. Cells were incubated with 10nM insulin for 20 min before glucose stimulation for 10 minutes (^3^H-2-deoxyglucose (50 nM; NET549250UC, PerkinElmer) diluted in non-radioactive glucose. Cells were washed with PBS and lysed with 0.1% Triton solution. Samples were transferred to scintillation vials containing scintillon and measured in a scintillation counter. Glucose uptake was normalized to protein concentration.

### GLUT4 translocation assay in murine brown adipocytes

For GLUT4 translocation experiments, immortalized murine brown adipocytes derived from previously characterized immortalized pre-brown adipocytes (Klein et al., 2002) were differentiated as described in previous sections. Differentiated brown adipocytes were electroporated with 45μg of HA-GLUT4-GFP. The next day the brown adipocytes were serum starved for 2 h and insulin was applied for 30 min. Cells were fixed with 3.7% formaldehyde for 6 minutes and GLUT4 on the cell surface was detected by an anti-HA antibody (1:1,000; MMS-101, BioLegend) without permeabilization. HA staining was visualized with Cy3-goat-anti-mouse (111-165-062, Jackson Immunoresearch). The surface HA signal was normalized to the GFP signal (total GLUT4). Perilipin, as an adipocyte differentiation marker was assessed using perilipin antibody (1:300; 9349, Cell Signaling Technology).

### Glucose and Insulin Tolerance tests

For Glucose Tolerance Tests (GTT), mice were fasted for 16 hours and acclimated to the procedure room with access to water. Mice were intraperitoneally (IP) injected with 2 g/kg of body weight glucose (G7528-1KG, Sigma). Blood was collected from the tail using Microvette^®^ CB 300 LH Heparin (16443100, SARSTEDT) and glucose levels were measured at 0, 3, 15, 30, 45, 60 time points. For the HFD experiments, 90 and 120 min time points were added. Plasma insulin levels were measured at 0, 3, 15 and 30 minutes after glucose injection. For Insulin Tolerance Tests (ITT), mice were fasted for 5 hours and acclimated to the procedure room with access to water. Mice were intraperitoneally (IP) injected with 0.5 unit/kg of body weight insulin for NCD-fed mice or 1 unit/kg of body weight insulin for HFD-fed mice insulin (Humulin R, Lilly) and glucose levels were measured at 0, 3, 15, 30, 45, 60 time points.

### Diet-induced obesity

For HFD studies, control and bR10KO mice were fed a HFD with 60% fat (D12492, Research Diets) or chow diet (D12450B, Research Diets) for 20 weeks (female mice) or for 25 weeks (male mice). The challenge with HFD started at 6 weeks of age. GTT was performed in females after 13 weeks on HFD and in males after 17 weeks on HFD.

### Cold-induced thermogenesis and thermoneutrality studies

Cold experiments were performed in an environmentally-controlled chamber with the temperature set at 4°C. Control and bR10KO mice where individually housed with *ad libitum* access to food and water for 14 days at 4°C. During this period, core body temperature was measured once a day using a rectal probe attached to a digital thermometer. After 14 days, mice were removed and immediately euthanized for tissue collection. Thermoneutrality experiments were performed at the Metabolic phenotyping center at Weill Cornell Medicine. Control and bR10KO mice were housed for 21 days in an environmentally-controlled chamber with the temperature set at 30°C. At day 19, the animals were subjected to a GTT: mice were fasted for 16 hours followed by an IP injection with 2 g/kg of body weight glucose (G7528, Sigma). After 21 days, core body temperature was measured using a rectal probe attached to a digital thermometer and mice were removed from the chamber and immediately euthanized for tissue collection.

### Plasma measurements

Plasma insulin (90082, Crystal Chem, Inc), and non-esterified fatty acids (NEFAs) (HR series, Wako) were measured following manufacturer instructions.

### Histology

For staining of adipose tissue for iBAT and iWAT samples were dissected and fixed in 10% Neutral buffer formalin overnight at 4°C. Samples were washed in PBS and processed for paraffin embedding using a TissueTek VIP 150 (8315-30-0016, Sakura) in the Electron Microscopy and Histology Core of WCM. Embedded blocks were sectioned at 5 μm thickness by the Laboratory of Comparative Pathology at Memorial Sloan Kettering. Slides were deparaffinized and stained with Hematoxylin & Eosin. The sections were imaged on a Zeiss AxioPlan microscope in the Electron Microscopy and Histology Core of WCM. Images were captured at 10x and 20x magnifications.

### RNA isolation and quantitative real-time polymerase chain reaction (qRT-PCR)

A piece of frozen tissue (BAT, iWAT or eWAT) was disrupted and homogenized in 1ml of Qiazol lysis reagent (79306, Qiagen) by using a Bead Mill homogenizer (Omni International). Samples were placed in the bench at room temperature for 5 minutes and 0.2ml of chloroform (C0549-1QT, Sigma) were added to each tube. The samples were vortexed and centrifuged at 12,000 rpm for 15 minutes. The aqueous phase was transferred in a new tube and mixed with 1 volume 70% ethanol. The volume was transferred to a RNeasy spin column and the protocol for the RNeasy mini kit (74106, Qiagen) was followed. Amplifications were done using primers summarized in Supplementary file 1, table 1.

### Protein extraction and western blot analysis

Protein was extracted from frozen tissue from eWAT, iWAT,BAT and liver by using cell lysis buffer (9803S, Cell Signaling) supplemented with complete protease and phosphatase inhibitor cocktail (78442, Thermo), and centrifuged at 12,000 rpm for 15 mins at 4°C. The supernatant was collected, and protein concentration was measured using a BCA protein quantification kit (Thermo Scientific, Waltham, MA). 35μg of protein was separated on 10% acrylamide gels and blotted onto nitrocellulose membranes. Membranes were blocked at room temperature for 1 h 5% milk in Tris-buffered saline-Tween (TBS-T). Membranes were incubated with primary antibodies (diluted 1:1000 in a 2% BSA-TBS-T) overnight at 4°C (anti-actin (AAN01-A, Cytoskeleton), anti-Rab10 (4262, Cell Signaling), anti-Fatty Acid Synthase (3180, Cell Signaling), anti-Phospho-Acetyl-Coa (11818, Cell Signaling) and anti-Acetyl-Coa Carboxylase (3676, Cell Signaling). Blots were incubated with goat anti–rabbit IgG-HRP (31460, Thermo Fisher) for 1 h at room temperature. The blots were scanned with the My ECL Imager (Thermo Fisher) quantified with Image J software.

### Quantification and statistical analysis

The Figure Legends indicate all of the subject numbers/replicates per study. Results are expressed as means ± SEM. All experiments were repeated at least three times. At least 4 mice per condition and per genotype were used as biological replicates for *in vivo* experiments. Data were analyzed using Prism 6.0 and Prism 8.0 (GraphPad), ImageJ and Metamorph softwares. Groups were compared with an analysis of variance (ANOVA) or a Student’s t test, and a P value < 0.05 was considered as significantly relevant.

## Notes

### Competing Interest Statement

The authors have declared no competing interest.

