## Supplemental Figures and Tables for "Blunting of insulin-stimulated glucose uptake in brown adipose tissue induces systemic metabolic dysregulation in female mice"

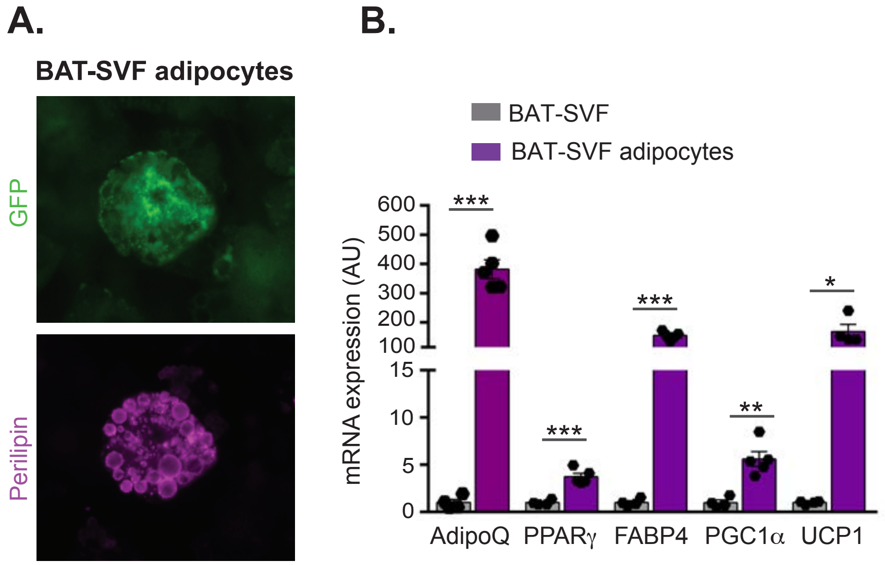


**Supplementary Figure 1. BAT-SVF adipocytes.** A. Immunofluorescence staining for perilipin in BAT-SVF adipocytes. B. qPCR analysis of mRNA levels of brown adipocyte markers of BAT-derived SVF cells before and after differentiation into adipocytes. *$p<0.05$, $p<0.001$, $p<0.0001$.

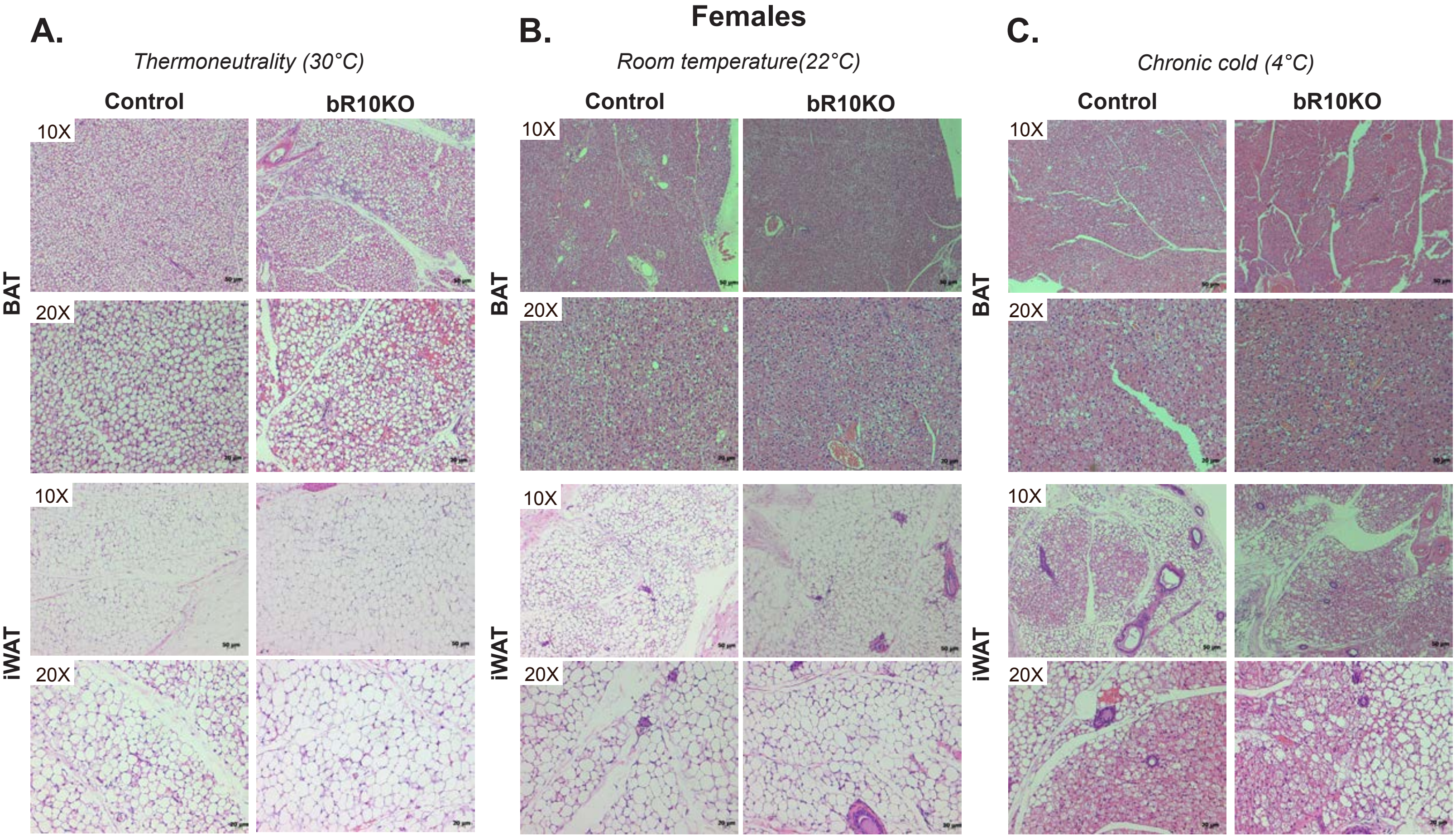


**Supplementary figure 2. Brown (BAT) and subcutaneous inguinal white adipose tissue (iWAT) from female mice.** A. BAT and iWAT from female mice acclimated to thermoneutrality for 21 days. B. BAT and iWAT from female mice housed at room temperature. C. BAT and iWAT from female mice acclimated to severe cold for 14 days. Images are shown at 10X and 20X magnification with 50μm and 20μm scale bars respectively.

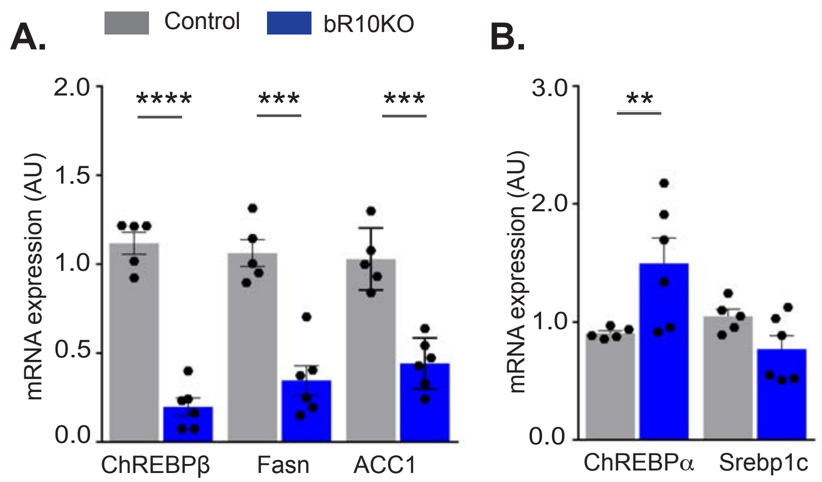


**Supplementary Figure 3. ChREBPβ and DNL-related gene expression are blunted in Rab10KO BAT from male mice.** mRNA expression for A. ChREBPβ, Fasn and ACC and B. ChREBPα and Srebp1c in BAT samples from 10 weeks of age thermoneutrality-acclimated male mice. **p <0.01, ***p<0.001 and ****p<0.0001.

**Supplementary Table 1.** Sequences for mouse primers used for qPCR.

| Species | Gene name | Primer |
|---|---|---|
| **Mouse** | **HPRT** | Forward: TGCTCGAGATGTCATGAAGG<br>Reverse: TATGTCCCCCGTTGACTGAT |
| **Mouse** | **ChREBPα** | Forward: CGACACTCACCCACCTCTTC<br>Reverse: TTGTTCAGCCGGATCTTGTC |
| **Mouse** | **ChREBPβ** | Forward: TCTGCAGATCGCGTGGAG<br>Reverse: CTTGTCCCGGCATAGCAAC |
| **Mouse** | **Srebp1c** | Forward: GGAGCCATGGATTGCACATT<br>Reverse: GGCCCGGGAAGTCACTGT |
| **Mouse** | **FASN** | Forward: GGAGGTGGTGATAGCCGGTAT<br>Reverse: TGGGTAATCCATAGAGCCCAG |
| **Mouse** | **DIO2** | Forward: AATTATGCCTCGGAGAAGACCG<br>Reverse: GGCAGTTGCCTAGTGAAAGGT |
| **Mouse** | **ACC1** | Forward: GATGAACCATCTCCGTTGGC<br>Reverse: GACCCAATTATGAATCGGGAGTG |
| **Mouse** | **UCP1** | Forward: AGGCTTCCAGTACCATTAGGT<br>Reverse: CTGAGTGAGGCAAAGCTGATTT |
| **Mouse** | **CIDEA** | Forward: TGACATTCATGGGATTGCAGAC<br>Reverse: GGCCAGTTGTGATGACTAAGAC |
| **Mouse** | **Adiponectin** | Forward: TGTTCCTCTTAATCCTGCCCA<br>Reverse: CCAACCTGCACAAGTTCCCTT |
| **Mouse** | **PRDM16** | Forward: CCAAGGCAAGGGCGAAGAA<br>Reverse: AGTCTGGTGGGATTGGAATGT |
| **Mouse** | **PPARγ** | Forward: GGAAAGACAACGGACAAATC<br>Reverse: TGGACACCATACTTGAGCAG |
| **Mouse** | **FABP4** | Forward: AAATGTGTGATGCCTTTGTG<br>Reverse: GTCACGCCTTTCATAACACA |
